## Supplementary figures and images for "Damage-induced basal epithelial cell migration modulates the spatial organization of redox signaling and sensory neuron regeneration"

### Supplemental Figure 1

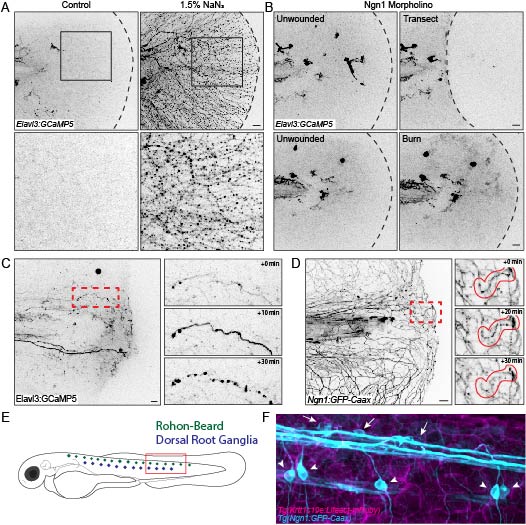

### Supplemental Figure 2

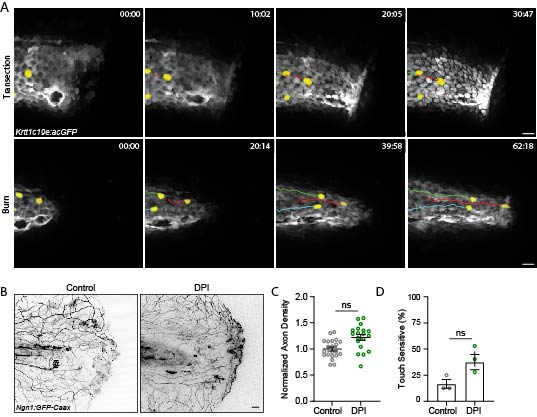

### Supplemental Figure 3

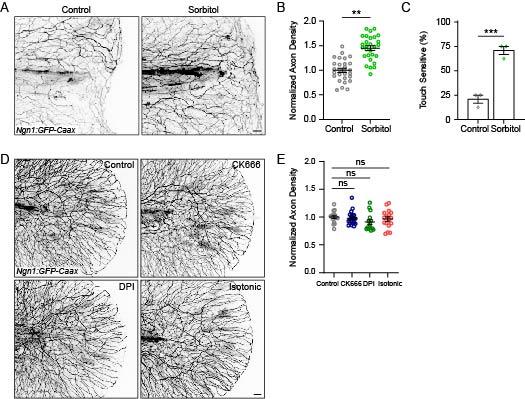
